# Out-of-Distribution Generalization from Labeled and Unlabeled Gene Expression Data for Drug Response Prediction

**DOI:** 10.1101/2021.05.25.445658

**Authors:** Hossein Sharifi-Noghabi, Parsa Alamzadeh Harjandi, Olga Zolotareva, Colin C. Collins, Martin Ester

## Abstract

Data discrepancy between preclinical and clinical datasets poses a major challenge for accurate drug response prediction based on gene expression data. Different methods of transfer learning have been proposed to address this data discrepancy. These methods generally use cell lines as source domains and patients, patient-derived xenografts, or other cell lines as target domains. However, they assume that they have access to the target domain during training or fine-tuning and they can only take labeled source domains as input. The former is a strong assumption that is not satisfied during deployment of these models in the clinic. The latter means these methods rely on labeled source domains which are of limited size. To avoid these assumptions, we formulate drug response prediction as an out-of-distribution generalization problem which does not assume that the target domain is accessible during training. Moreover, to exploit unlabeled source domain data, which tends to be much more plentiful than labeled data, we adopt a semi-supervised approach. We propose Velodrome, a semi-supervised method of out-of-distribution generalization that takes labeled and unlabeled data from different resources as input and makes generalizable predictions. Velodrome achieves this goal by introducing an objective function that combines a supervised loss for accurate prediction, an alignment loss for generalization, and a consistency loss to incorporate unlabeled samples. Our experimental results demonstrate that Velodrome outperforms state-of-the-art pharmacogenomics and transfer learning baselines on cell lines, patient-derived xenografts, and patients. Finally, we showed that Velodrome models generalize to different tissue types that were well-represented, under-represented, or completely absent in the training data. Overall, our results suggest that Velodrome may guide precision oncology more accurately.

## Introduction

The goal of drug response prediction based on the genomic profile of a patient (also known as pharmacogenomics) -- a crucial task of precision oncology -- is to utilize the omics features of a patient to predict response to a given drug (Garraway, Verweij, and Ballman 2013; Cronin et al. 2018; Marquart, Chen, and Prasad 2018; Pal et al. 2019). Unfortunately, patient datasets with drug response are often small or not publicly available which motivated the creation of large-scale preclinical resources such as patient-derived xenografts (PDX) (Gao et al. 2015) or cancer cell lines (Garnett et al. 2012; Barretina et al. 2012; Basu et al. 2013; Seashore-Ludlow et al. 2015; Klijn et al. 2015; Iorio et al. 2016; Haverty et al. 2016) as proxies for patients.

Although preclinical datasets are viable proxies for patients, they differ in important ways from patients due to basic biological differences such as the lack of tumor microenvironment/the immune system (Mourragui et al. 2019; Hossein Sharifi-Noghabi, Peng, et al. 2020) -- This has been a source of discussions in the community showing discrepancy between preclinical resources (Haibe-Kains et al. 2013; Safikhani et al. 2016) and providing evidence regarding consistency between them (Haverty et al. 2016; Mpindi et al. 2016; Geeleher et al. 2016).

Transfer learning has emerged as a machine learning paradigm for such scenarios (Pan and Yang 2010; Neyshabur, Sedghi, and Zhang 2020), where we have access to different datasets from multiple resources (known as source domains) and want to make predictions for a dataset of interest (known as target domain) and it has been employed in different problems (Taroni et al. 2019; Raghu et al. 2019; Holmberg et al. 2020; Hu et al. 2020). Various methods of transfer learning have been proposed in the context of drug response prediction. These methods either address these discrepancies implicitly (Hossein Sharifi-Noghabi et al. 2019; Snow et al. 2020; Kuenzi et al. 2020), or explicitly which means they assume that the model has access to the desired labeled or unlabeled target domain during training (Hossein Sharifi-Noghabi, Peng, et al. 2020; Mourragui et al. 2019, 2020; Ma et al. 2021; Zhu et al. 2020; Salvadores, Fuster-Tormo, and Supek 2020; Najgebauer et al. 2020; Peres da Silva, Suphavilai, and Nagarajan 2021; Warren et al. 2021).

However, in the real-world we do not have access to the target domain(s) during training the model on the source domain, e.g., we do not know future patients that may walk into a clinic. Nevertheless, the trained model should generalize to the target domain and be able to make predictions for samples encountered during the deployment time. Since generating large high-quality labeled preclinical datasets is an expensive and time-consuming process and we do not know response to a given drug in the target domain (e.g., future patients), there is a need for a computational method that takes not only labeled but also unlabeled source domain data as input and learns a representation that generalizes to a future target domain. This problem is known as out-of-distribution generalization or domain generalization, where the target domain is not accessible during training (Gulrajani and Lopez-Paz 2020; J. Wang et al. 2021; Zhou et al. 2021). Out-of-distribution generalization is particularly important for biomedical applications (Zhang et al. 2021).

There are two main approaches to out-of-distribution generalization: 1) generalizing via learning domain-invariant features (J. Wang et al. 2021), and 2) generalizing via learning hypothesis-invariant features (Zhao et al. 2020; Z. Wang, Loog, and van Gemert 2021). In the first approach, the goal is to map the input domains to a shared feature space in which the features of all domains are aligned, i.e. look similar to each other. However, forcing different domains to have very similar features is not always feasible because different domains may have unique characteristics, and completely aligning them ignores these unique characteristics. The second approach does not align the features but rather the predictions across domains. The idea is that if the extracted features of input domains are similar enough for an accurate predictor to make similar predictions, forcing the features to be more similar is not required anymore. We note that there is no existing method for out-of-distribution generalization applicable to both regression and classification (Hossein Sharifi-Noghabi, Asghari, et al. 2020), for either of the two approaches, that can exploit both labeled and unlabeled source domains.

In this paper, we propose Velodrome, a deep neural network method that combines the two above approaches and exploits both labeled and unlabeled samples. Velodrome takes gene expression from cell line (labeled) and patient (unlabeled) datasets as input domains and predicts the drug response (measured as area above dose-response curve, AAC) via a shared (between cell lines and patients) feature extractor and domain-specific predictors. The feature extractor and the predictors are trained using a novel loss function with three components: 1) a standard supervised loss to make the features predictive of drug response, 2) a consistency loss to exploit unlabeled samples in learning the feature representation, and 3) an alignment loss to make the features generalizable. We designed the loss function to balance between learning domain-invariant and hypothesis-invariant features. To the best of our knowledge, Velodrome is the first method for semi-supervised out-of-distribution generalization from labeled cell lines and unlabeled patients to different preclinical and clinical datasets.

We evaluated the performance of Velodrome and state-of-the-art methods of supervised out-of-distribution generalization, domain adaptation, and semi-supervised learning in terms of a diverse range of metrics including Pearson and Spearman correlation, the Area Under the Receiver Operating Characteristic curve (AUROC), and the Area Under the Precision-Recall curve (AUPR). We observed that Velodrome achieved substantially better performance across different clinical and preclinical pharmacogenomics datasets for multiple drugs, demonstrating the potential of semi-supervised out-of-distribution generalization for drug response prediction, a crucial task of precision oncology. Moreover, we showed that the responses predicted by Velodrome for TCGA patients (unlabeled, i.e. without drug response) with prostate and kidney cancers had statistically significant associations with the expression values of the target genes of the studied drugs. This shows that Velodrome captures biological aspects of drug response. Finally, although Velodrome was trained only on solid tissue types, we showed that it made accurate predictions for cell lines originating from non-solid tissue types, showcasing the out-of-distribution capabilities of the Velodrome model.

## Results

### Datasets

We employed the following resources throughout this paper:

1. Patients without drug response: more than 3,000 samples obtained from TCGA (Cancer Genome Atlas Research Network et al. 2013) breast (TCGA-BRCA), lung (TCGA-LUAD), pancreatic (TCGA-PAAD), kidney (TCGA-KIRC), prostate (TCGA-PRAD), Myeloid (TCGA-LAML), and lymphoma (TCGA-DLBC) cohorts with RNA-seq data.
2. Cell lines with drug response: The Cancer Therapeutics Response Portal (CTRPv2) (Basu et al. 2013; Seashore-Ludlow et al. 2015), The Genomics of Drug Sensitivity in Cancer (GDSCv2) (Garnett et al. 2012; Iorio et al. 2016), and The Genentech Cell Line Screening Initiative (gCSI) (Haverty et al. 2016; Klijn et al. 2015) pan-cancer datasets with a total of more than 2000 samples with RNA-seq data and AAC as the measure of the drug response across 11 drugs (in common for the three datasets). We focused on the following drugs for this paper: Erlotinib, Docetaxel, Paclitaxel, and Gemcitabine.
3. PDX samples with drug response: PDX Encyclopedia (PDXE) dataset (Gao et al. 2015) is a collection of more than 300 PDX samples with RNA-seq data screened with 34 drugs. We use the reported measure of response in RECIST (Schwartz et al. 2016) for Gemcitabine, Erlotinib, and Paclitaxel obtained from supplementary material of (Gao et al. 2015).
4. Patients with drug response: 2 cancer-specific datasets with microarray data and RECIST as the measure of drug response for Docetaxel (Hatzis et al. 2011), Paclitaxel (Hatzis et al. 2011), and Erlotinib (Byers et al. 2013). Plus, a pan-cancer dataset obtained from TCGA patients treated with Gemcitabine (Ding, Zu, and Gu 2016). We use clinical annotations of the drug response for some patients which were obtained from supplementary material of (Ding, Zu, and Gu 2016).

Supplementary Table 1, 2, and 3 present characteristics of these datasets and indicate whether they were used as source domain for training or target domain for test.

### Velodrome Overview

The proposed Velodrome method takes gene expression and AAC of cell line datasets (CTRPv2 and GDSCv2) as well as gene expression of patients without drug response (TCGA dataset) and learns a predictive and generalizable representation. To achieve this, Velodrome employs a shared feature extractor, which takes the gene expression of CTRPv2 and GDSCv2 samples and maps them to a shared feature space, and domain-specific predictors (e.g. one for CTRPv2 and one for GDSCv2), which take the feature representation of the gene expression and predict the drug response.

The parameters are optimized using a novel objective function consisting of three loss components. 1) a standard supervised loss to make the representation predictive of drug response, 2) a consistency loss to exploit unlabeled samples in learning the representation, and 3) an alignment loss to make the representation generalizable.

The idea of the standard supervised loss is to make the representation predictive of the drug response via a mean squared loss.

To incorporate unlabeled patient samples, we add a consistency loss. The idea is to first extract features from patient samples using the feature extractor and then assign pseudo-labels to them by utilizing the predictors associated with CTRPv2 and GDSCv2. The consistency loss takes the pseudo-labels (i.e., predictions) from the predictors and regularizes the parameters of the feature extractor and the predictors by the *l*_2_ distance between the predictions of CTRPv2 predictor and those of the GDSCv2 predictor.

Finally, to make the feature representation generalizable, we add an alignment loss that regularizes the parameters of the feature extractor. This alignment loss takes the extracted features of any two input domains (eg., CTRPv2 and TCGA or CTRPv2 and GDSCv2) and minimizes the difference between the covariance matrices of those domains.

Figure 1 illustrates the schematic overview of the Velodrome method.

**Figure 1.**
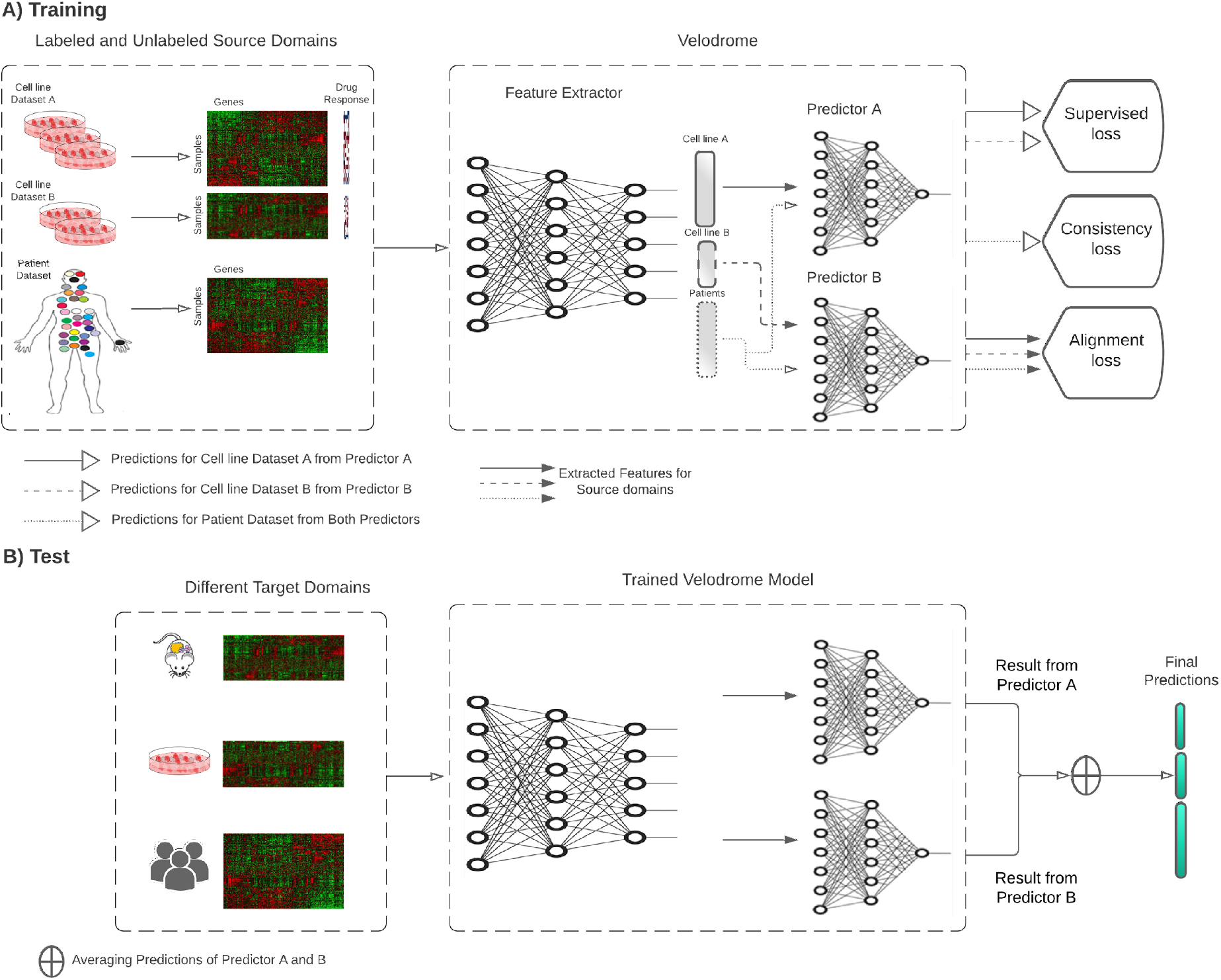
The schematic overview of the Velodrome method with three source domains (two labeled and one unlabeled). A) At training time, the feature extractor receives data from different source domains and extracts high-level abstract features. The extracted features of each labeled domain (cell line dataset) are input to the corresponding domain-specific predictor. Predictions are used to optimize the parameters of the predictors and the feature extractor via a standard supervised loss function. The extracted features of the unlabeled domain (patient dataset) are input to both predictors, and the predictions are used to optimize the parameters of predictors and the feature extractor via a consistency loss function. The extracted features of all source domains are used to optimize the parameters of the feature extractor via an alignment loss function. B) At test time, the trained Velodrome model receives samples from different target domains, extracts features and makes predictions using the trained predictors. The predictions are then averaged to generate the final predictions for each sample.

### Evaluation

Drug response prediction using multiple labeled and unlabeled domains can be viewed in three approaches: 1) under the assumption that there is no data discrepancy, it can be viewed as a semi-supervised learning problem, 2) under the assumption that unlabeled patient samples are proxies to future patients, it can be viewed as an unsupervised domain adaptation problem, and 3) under the assumption that a generalizable representation can be obtained via only labeled domains, it can be viewed as a supervised domain generalization problem. It is important to note that the main contribution of the Velodrome method is that it is the first semi-supervised domain generalization method for drug response prediction.

To evaluate the performance of Velodrome, we compared it against the state-of-the-art methods of each approach. For the first approach, we compared Velodrome to Mean Teacher (Tarvainen and Valpola 2017) which is the state-of-the-art deep neural network for semi-supervised learning (Yang and Xu 2020). For The second approach, we compared Velodrome to PRECISE as a non-deep learning method based on subspace alignment and (Saito et al. 2018) as a deep learning method based on adversarial domain adaptation via disagreement between predictors. Finally, for the third approach, we compared Velodrome to Ridge-ERM (Ridge Regression) as a non-deep learning baseline and DeepAll-ERM as a deep learning baseline. Both of them are categorized as methods of Empirical Risk Minimization (ERM). In an extensive benchmark in the context of computer vision, ERM methods tended to achieve state-of-the-art performance for out-of-distribution generalization (Gulrajani and Lopez-Paz 2020). They are trained in a supervised fashion by merging all available labeled input domains.

### Velodrome makes accurate predictions for cell lines

To investigate the generalization of Velodrome to other cell line datasets, we employed the gCSI dataset as the target domain and reported the performance of Velodrome and the baselines in terms of the Pearson and the Spearman correlation on this dataset. On average±sd over all drugs, DeepAll-ERM achieved the best performance (0. 52 ± 0. 09 for Pearson correlation coefficient and 0. 48 ± 0. 09for Spearman correlation coefficient - Figure 2D). Velodrome achieved the second best performance (0. 48 ± 0. 09 for Pearson correlation coefficient and 0. 45 ± 0. 07for Spearman correlation coefficient - Figure 2A and D). Ridge-ERM (0. 46 ± 0. 07-Figure 2A and D) and Mean Teacher (0. 43 ± 0. 07-Figure 2A and D) had the third best performance in terms of Pearson and Spearman correlation, respectively. These results indicate that although Velodrome is not the best performing model, it is fairly competitive on cell lines and generalizes well (Figure 2A).

**Figure 2.**
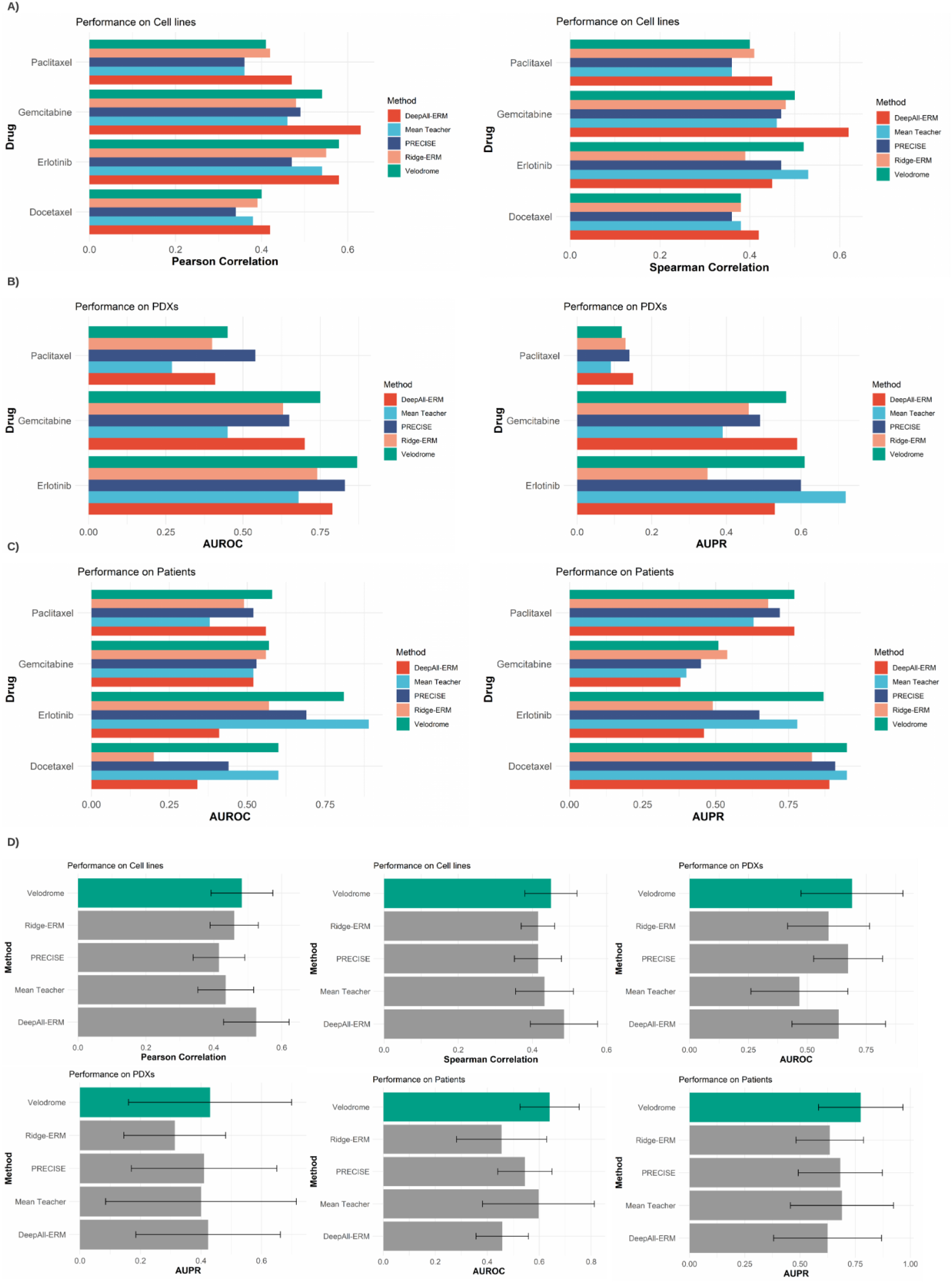
Comparison of Velodrome and state-of-the-art of drug response prediction methods on: (A) cell lines in terms of Pearson and Spearman correlations, (B) PDX models in terms of the Area Under the Receiver Operating Characteristic curve (AUROC) and the Area Under the Precision-Recall curve (AUPR), (C) patients in terms of AUROC and AUPR. (D) average±sd over the studied drugs for each method, Velodrome has the best or the second best performance on cell lines, PDX models, and patients compared to the baseline methods. The error bar indicates standard deviation of performance across studied drugs..

### Velodrome makes accurate predictions for PDXs samples

To investigate generalization of Velodrome to PDX samples, we employed the PDXE dataset as the target domain and reported the performance of Velodrome and the baselines discussed above in terms of the AUROC and the AUPR. On average±sd over all drugs,, Velodrome achieved the best performance compared to the baselines (for 0. 69 ± 0. 21in AUROC and 0. 43 ± 0. 26in AUPR-Figure 2B and D). PRECISE obtained the second best performance in terms of AUROC (0. 67 ± 0. 14- Figure 2B and D) and DeepAll-ERM in terms of AUPR (0. 42 ± 0. 23- Figure 2B and D). Similarly, DeepAll had the third best performance in terms of AUROC (0. 63 ± 0. 19-Figure 2B and D) and PRECISE had the third best performance in terms of AUPR (0. 41 ± 0. 24-Figure 2B and D). These results indicate that utilizing both labeled and unlabeled samples from cell lines and patients improves drug response prediction on PDX samples.

### Velodrome makes accurate predictions for patients

To investigate the generalization of Velodrome to patient samples, we employed the patient datasets obtained from clinical trials as target domains and reported the performance of Velodrome and the baselines discussed above in terms of AUROC and AUPR. On average±sd over all drugs, Velodrome achieved the best performance compared to the baselines and outperformed the competitors (0. 64 ± 0. 11in AUROC and 0. 77 ± 0. 19in AUPR-Figure 2C-D). Mean Teacher obtained the second best performance (0. 59 ± 0. 21in AUROC and 0. 69 ± 0. 23in AUPR-Figure 2C-D) and PRECISE had the third best performance (0. 54 ± 0. 1in AUROC and 0. 68 ± 0. 18in AUPR-Figure 2C-D). Interestingly, these three top-performing methods take for inputs both labelled and unlabeled samples, in contrast to other baselines considering only labeled samples. These results indicate that incorporating unlabeled patient data along with labeled data significantly improves the generalization performance on patients. However, the results also demonstrate the advantage of learning features that are domain-invariant and hypothesis-invariant for out-of-distribution generalization, because the PRECISE method only ensures a domain-invariant representation.

### Velodrome outperforms the baselines over multiple independent runs

To maximize the reproducibility, we utilized a fixed random seed for all methods (Velodrome and the baselines) and found the best settings for the hyper-parameters of each method with that seed. To investigate the performance of the best trained Velodrome model for each drug and those of the baselines, we re-trained all of the models from scratch using the same settings with 10 different random seeds and reported mean±sd for each method (Figure 3A) over all runs and studies drugs. Although we observed that the average performance (over the studied drugs) of all methods decreased, Velodrome still achieved the best performance on patients in terms of both AUROC and AUPR, and also the best performance in terms of both Pearson and Spearman correlation on cell lines. PRECISE and DeepAll-ERM obtained the best performance on PDX samples in terms of AUROC and AUPR, respectively (the performance of these two methods tied on AUPR). Velodrome had the third best performance in terms of AUROC and AUPR on PDX samples. Overall, these results indicate that Velodrome is more accurate and competitive compared to baselines particularly on patients and cell lines.

**Figure 3.**
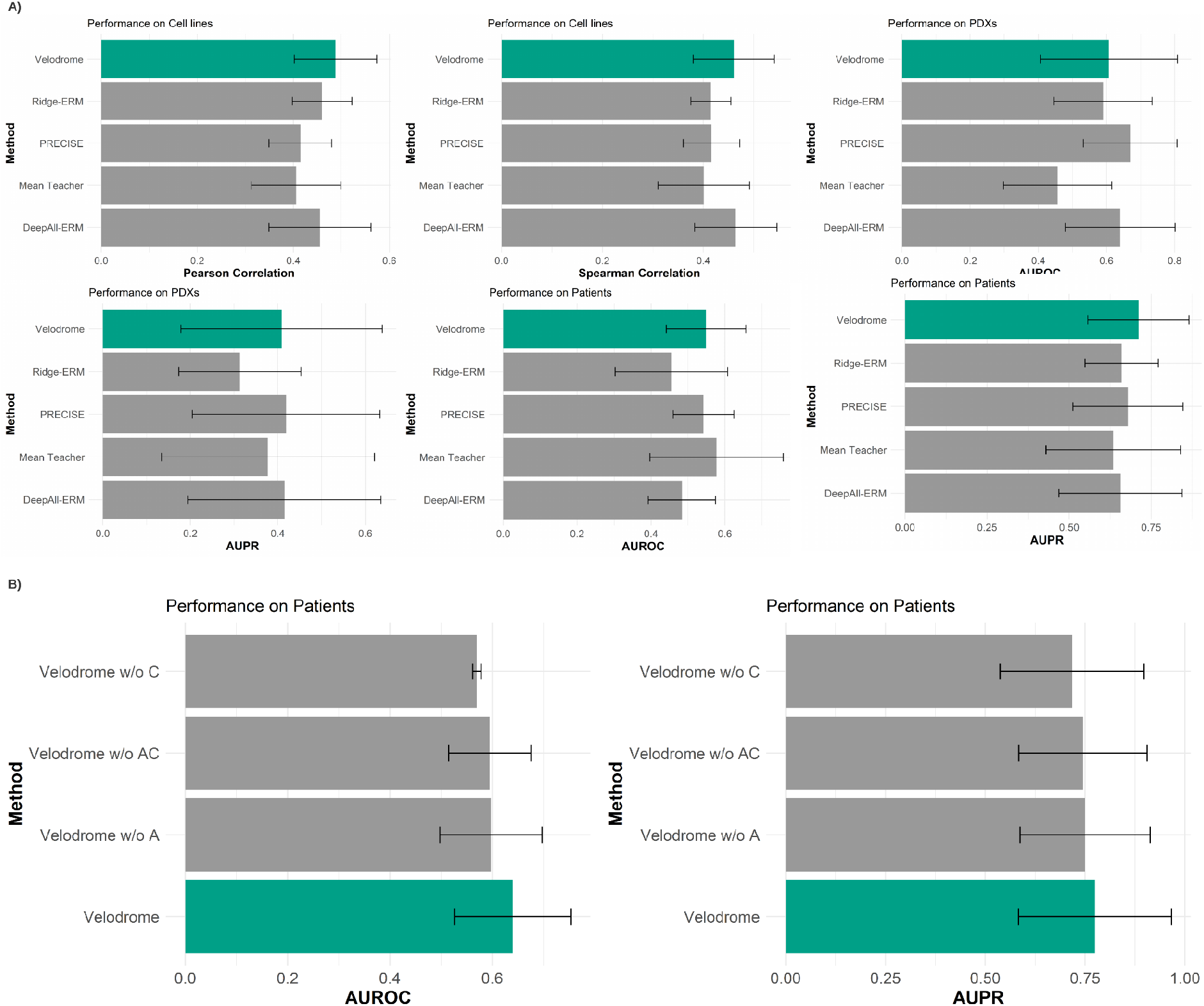
(A) Comparison of the average±sd performance of the Velodrome and the state-of-the-art of drug response prediction methods over 10 independent runs for the studied drugs in terms of Pearson and Spearman correlations as well as the Area Under the Receiver Operating Characteristic curve (AUROC) and the Area Under the Precision-Recall curve (AUPR). (B) average±sd of an ablation study to examine different components of the Velodrome on patients in terms of AUROC and AUPR. The error bar indicates standard deviation of performance across studied drugs.

### The complete version of Velodrome demonstrates the best performance

We performed an ablation study to investigate the impact of the different loss components of Velodrome separately. We studied three scenarios as follows: “*Velodrome w/o A”* represents a version of Velodrome without the alignment loss component, which means the neural network only uses supervised and semi-supervised losses. “*Velodrome w/o C”* represents a version of Velodrome without the consistency loss, which means the neural network only considers the supervised loss and the alignment loss. Finally, “*Velodrome w/o AC”* represents a version of Velodrome without both the alignment and the consistency loss, which means the neural network employed only has a standard supervised loss. Our results on patients demonstrate that on average±sd (over all drugs for 10 independent runs), the complete version of Velodrome outperforms its variants which indicates the added value of both alignment and consistency losses (Figure 3-B). Interestingly, removing the consistency loss from the objective function had the biggest impact on the Velodrome performance on patients. This may suggest that hypothesis alignment plays a more critical role than feature alignment for out-of-distribution generalization, which is compatible with recent observations in computer vision applications (Zhao et al. 2020).

### Velodrome generalizes to well-represented tissue types

To evaluate the performance of Velodrome on patients, we followed the experimental design of previous pharmacogenomics methods and designed an association study based on the known associated target genes for the investigated drugs (Geeleher et al. 2017; Mourragui et al. 2019; Hossein Sharifi-Noghabi et al. 2019; Hossein Sharifi-Noghabi, Peng, et al. 2020). In this analysis, we employed the TCGA Kidney cancer cohort (TCGA-KIRC) as a tissue type well represented in our cell line datasets. In GDSCv2 and CTRPv2 combined, more than 3.3% of the samples originated from this tissue type (Supplementary Figure 1).

We trained Velodrome models for each drug (Docetaxel, Erlotinib, Paclitaxel, and Gemcitabine) and applied them to the gene expression data of the patients of this cohort to predict their response. Then, we fit a linear regression model to the level of expression of the known target genes of these drugs and the responses predicted by Velodrome. Based on the corrected p-values (two-tailed t-test) obtained from this multiple linear regression using the bonferroni correction method, there are a number of statistically significant associations between the target genes of the studied drugs and the responses predicted by Velodrome. For Docetaxel, MAP2 had a statistically significant association(*P* < 10^−6^). For Erlotinib, EGFR and ERBB2 had statistically significant associations (both *P* < 10^−6^). For Paclitaxel BCL2 and MAP2 had significant associations (both *P* < 10^−6^). Finally, for Gemcitabine, CMPK1 demonstrated a significant association(*P* < 10^−6^). These results suggest that the responses predicted by Velodrome are not random but capture biological aspects of the drug response. BCL2 was not significant (*P* > 0. 05) for Docetaxel after hypothesis correction at level α = 0. 05.

### Velodrome generalizes to under-represented tissue types

To further evaluate the performance of Velodrome, we performed a similar association study on the prostate cancer cohort in TCGA (TCGA-PRAD). We chose prostate because unlike kidney, prostate is a tissue type under-represented in our cell line datasets (only 0.3% of the samples originated from this tissue).

Similar to TCGA-KIRC, the Velodrome predictions for TCGA-PRAD patients demonstrated significant associations with known target genes of the studied drugs. For Docetaxel, MAP2 showed a statistically significant association (*P* < 10^−6^). For Erlotinib, both EGFR and ERBB2 showed statistically significant associations (both *P* < 10^−6^). For Paclitaxel, BCL2 (*P* = 8 × 10^−6^) and MAP2 (*P* < 10^−4^) had significant associations. Finally, for Gemcitabine, CMPK1 demonstrated significant association (*P* < 10^−6^). These results confirm again that the responses predicted by Velodrome are not random and they capture biological aspects of the drug response even for a tissue under-represented in the source domain. In this cohort, BCL2 was not significant (*P* > 0. 05) for Docetaxel after hypothesis correction at level α = 0. 05.

### Velodrome generalizes to new tissue types

Finally, we trained Velodrome and the baseline methods only on samples (cell lines and patients) that originated from solid tissue types because non-solid tissues such as haematopoietic and lymphoid have different molecular and pharmacological profiles compared to solid ones (Noghabi et al. 2021). Therefore, we wanted to examine the out-of-distribution capability of the Velodrome models on these tissue types that were completely absent during training. For that, we tested the trained Velodrome models for the studied drugs on samples originated from non-solid tissues in the gCSI cell line dataset and evaluated the performance in terms of Pearson correlation between the predictions and the actual AAC values and reported two-tailed p-value as well.

For Erlotinib and Gemcitabine, Velodrome demonstrated significant correlations of 0.4 (*P* = 5 × 10^−3^) and 0.39 (*P* = 4 × 10^−3^), respectively. For Docetaxel and Paclitaxel, Velodrome did not make accurate predictions and had poor correlations of −0.07 and −0.02, respectively (both *P* > 0. 05).

As a baseline to compare the Velodrome performance on non-solid tissues, we trained a Ridge Regression model on samples originated from non-solid tissues in CTRPv2 and GDSCv2 datasets and tested this predictor on non-solid samples of gCSI dataset. Therefore, we built a predictor specifically for non-solid samples and the performance of this model should act as an upper bound for the Velodrome. Similar to the Velodrome results, this predictor also achieved significant correlations of 0.34 (*P* = 10^−2^) and 0.39 (*P* = 5 × 10^−3^) for Erlotinib and Gemcitabine and negative correlations of −0.11 (*P* > 0. 05) and −0.4 (*P* = 4 × 10^−3^) for Docetaxel and Paclitaxel, respectively. These results suggest that Velodrome is as accurate (and even more accurate in the case of Erlotinib) as a non-solid predictor on these tissues even though it did not utilize them during training. The poor/negative correlation for Docetaxel and Paclitaxel may be dataset specific, particularly in the case of Paclitaxel where the non-solid predictor had a significant negative correlation, and requires further study.

### Within-tissue generalization

We expanded on the within-tissue performance of Velodrome to further study the generalization capability of the trained models. We compared Velodrome predictions with the AAC baseline correlation in the utilized datasets. To obtain this baseline correlation, we selected cell lines with solid tissues that are common between our training data (cell lines in CTRPv2 and GDSCv2) and our test data (cell lines in gCSI). This baseline correlation indicates how much train and test data agree with each other in terms of drug response (AAC). We expect to see a comparable correlation for Velodrome predictions if the given model is accurate enough. Our results demonstrated that for the majority of the drugs, Velodrome achieved comparable performance compared to the baseline correlation in terms of Pearson and Spearman correlations (Figure 4). This reconfirms our previous results on generalization within solid tissues.

**Figure 4.**
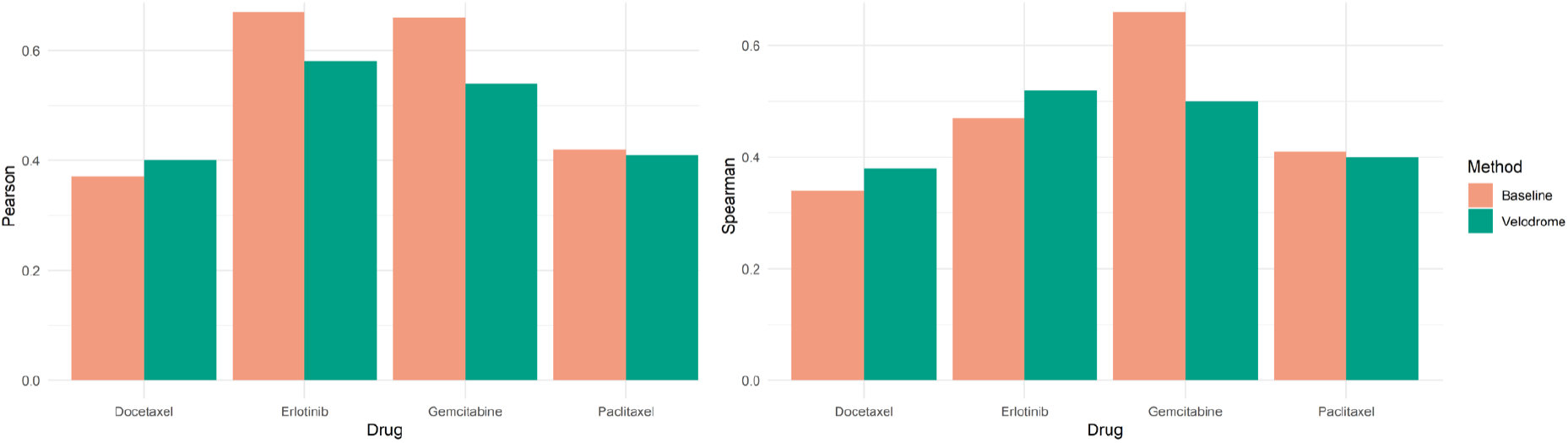
Comparison of Velodrome predictions to the baseline correlation in terms of Pearson (left side) and Spearman correlation (right side). The baseline correlation is obtained by calculating the correlation of cell lines in common between train and test data for each of the studied drugs. This baseline correlation shows how much train and test data are consistent in terms of Area Above dose-response Curve (AAC).

### Cross-tissue generalization

We also expanded the cross-tissue generalization analysis by repeating a similar association study of well- and under-represented tissues but this time for two TCGA cohorts with non-solid tissue types. For that, we obtained TCGA-LAML and TCGA-DLBC datasets for Acute Myeloid Leukemia and Diffuse large B-cell lymphoma, respectively. Similar to TCGA-KIRC and TCGA-PRAD, Velodrome predictions for TCGA-LAML and TCGA-DLBC patients demonstrated significant associations with known target genes of the studied drugs.

For Docetaxel, MAP2 showed a statistically significant association in TCGA-LAML (*P* = 10^−5^) and TCGA-DLBC (*P* = 0. 02) but BCL2 was not significant in these cohorts. For Erlotinib, EGFR and ERBB2 showed statistically significant associations for TCGA-LAML (both *P* < 10^−6^) and these two genes were also significant in TCGA-DLBC (8 × 10^−6^ and 4 × 10^−7^ respectively). for Gemcitabine, CMPK1 demonstrated significant association (*P* < 10^−6^) in both cohorts. Finally, for Paclitaxel, only BCL2 (*P* = 4 × 10^−4^) had significant associations for patients in the TCGA-LAML cohort and we did not observe any significant associations for TCGA-DLBC. Our results demonstrate that for non-solid patient data, associations were not significant for Paclitaxel which align with the previous results where Velodrome did not generalize to non-solid cell lines for Paclitaxel. For Docetaxel, p-values are larger than what we observed for solid tumors which again aligns the previous results where Velodrome did not generalize to non-solid cell lines. However, we observed significant associations for Erlotinib and Gemcitabine which is consistent with Velodrome accurate performance on non-solid cell lines. Overall, these new results also reconfirm cross-tissue generalization of Velodrome models.

## Discussion

From the biological point of view, we found interesting connections between the known target genes of the studied drugs and the TCGA cohorts that we investigated (TCGA-PRAD and TCGA-KIRC). For example, BCL2 has known connections to prostate cancer progression (Chaudhary, Abel, and Lalani 1999; Catz and Johnson 2003) and survival (Renner et al. 2017). More importantly, the expression of BCL2 may have an antiapoptotic activity against androgen which is a key player in prostate cancer (Chaudhary, Abel, and Lalani 1999). Similarly, BCL2 can also act as an oncoprotein in kidney cancer (Paraf et al. 1995) and therapeutics roles (Adams and Cory 2007; Delbridge et al. 2016).

As another example, Microtubule-Associated Proteins including MAP2 have also been associated with different cancers including prostate (Bhat and Setaluri 2007) and kidney cancers (He et al. 2020). Moreover, Microtubule-targeting chemotherapy agents, Docetaxel and Paclitaxel, have been used in combination with anti-androgen therapeutics to increase the survival rate in prostate cancer patients (Martin, Kamelgarn, and Kyprianou 2014). Prostate cancer progression and lethal outcome have been associated with metabolic signaling pathways and CMPK1 (it mediates the mechanism of action for Gemcitabine) was shown to be highly expressed in prostate cancer patients (Kelly et al. 2016). A combination of Gemcitabine and other chemotherapy agents has shown to be effective for a subtype of kidney cancer (Numakura et al. 2014). Finally, EGFR and ERBB2 have been associated with different cancer types including prostate (El Sheikh et al. 2004; Pignon et al. 2009) and kidney (Reid et al. 2007) and they both showed therapeutic opportunities and increase in survival (Gordon et al. 2009).

From the computational point of view, it has been shown that methods of empirical risk minimization (ERM) are highly competitive for supervised domain generalization (Gulrajani and Lopez-Paz 2020). Therefore, it was also expected to see a competitive performance for a semi-supervised method (Mean Teacher) for the semi-supervised domain generalization setting. Moreover, Velodrome, PRECISE, and Mean Teacher were designed to take both labeled and unlabeled samples and, therefore, were expected to achieve better performance on patients than DeepAll-ERM and Ridge-ERM. On the other hand, these two methods achieved better performance on cell lines which makes sense since they were trained on cell lines.

We considered only TCGA-BRCA, TCGA-PAAD, and TCGA-LUAD for training, because these tissue types were well-represented in our cell line datasets (Supplementary Figure 1) and because the four studied drugs were treatment options for these cell lines. This selection increases the relevancy of labeled (cell lines) and unlabeled (TCGA patients) data. Relevancy has been shown to improve semi-supervised learning performance even when both labeled and unlabeled datasets are imbalanced (Yang and Xu 2020), which is the case for drug response prediction.

Although methods of adversarial domain adaptation have shown great performance in different applications, especially computer vision (Ganin and Lempitsky 2015; Tzeng et al. 2017; Chen et al. 2017), we did not consider them as baselines because they were clearly outperformed by PRECISE (which we do use as baseline) in a recent study (Hossein Sharifi-Noghabi, Peng, et al. 2020).

Although gene expression data has been shown many times to be the most effective genomic data type for drug response prediction (Iorio et al. 2016; Costello et al. 2014), in principle Velodrome can be extended to incorporate other omics data types. Especially promising are proteomics data (Ali et al. 2018) and germline variants (Menden et al. 2018), due to their predictive power. The advantage of proteomics is that it is closer to the phenotype and gene expression and protein abundance can be quite discordant. Velodrome can also be extended to incorporate additional information about the drug, such as the chemical representation, to improve the performance (Jiang et al. 2020). Finally, we did not discuss the explainability of the Velodrome model, but we note that the feature extractor of Velodrome can be replaced by a knowledge-based network (Snow et al. 2020) to offer explainability and transparency (Yu et al. 2018). A major limitation of our work is the output space discrepancy between cell lines, PDX samples, and patients, because on cell lines the drug response is measured based on the concentration of the drug but on PDX samples and patients the response is measured based on the change in the tumor volume after treatment. A recent method adjusts for this output space discrepancy and improves the prediction performance (Hossein Sharifi-Noghabi, Peng, et al. 2020), but this method requires access to the target domain during training which violates the assumption of out-of-distribution generalization. In this work, we used AAC as the measure of drug response in cell line datasets and treated it as a score for making predictions for patients and PDX samples. However, measuring AAC is dependent on the tested concentration range which generally differs between different pharmacogenomics studies. Recent efforts have demonstrated that adjusting concentration ranges across different datasets improves the prediction performance (Pozdeyev et al. 2016; Xia et al. 2021), however, we did not consider this adjustment because it reduces the sample size substantially.

Although drug response prediction was the driving problem of the Velodrome method, in principle it is also applicable to other problems, especially in clinical settings. For example, prostate cancer has steady progression, meaning datasets for this cancer have a limited number of labeled samples with respect to long-term outcomes such as metastasis but more unlabeled samples (H. Sharifi-Noghabi et al. 2019). This makes Velodrome applicable to metastasis prediction via labeled and unlabeled gene expression data. Another example is the case of rare cancers for which obtaining large training datasets is extremely difficult. Velodrome demonstrated cross-tissue generalization which makes it applicable for these rare tissue types. Both of these cases can be promising future directions.

Indeed, our results suggest that owing to its ability for cross-tissue generalization, Velodrome has a potential to predict the response even for cancer types under-represented or completely absent in the training data. The cross-tissue generalization is particularly interesting because some previous studies have shown basic differences in gene expression between solid and non-solid tissues (Torrente et al. 2016; Lukk et al. 2010), however, our empirical results demonstrated that Velodrome models trained on solid tissues generalize to non-solid tissues in cell lines and patients. This may suggest shared component(s) in basal transcription factors. One example is cyclin-dependent kinases (CDKs) which are targets for global transcription inhibition in numerous drugs (Villicaña, Cruz, and Zurita 2014). Interestingly, some of these drugs are treatments for both solid and non-solid cancers.

In addition, another explanation for cross-tissue generalization can be the presence of drivers shared across cancer types. Different studies have identified pan-cancer driver genes across solid and non-solid tissues such as TP53, ERBB2, EGFR, or EML4-ALK (Mano 2008; Zapata et al. 2017; Abou Dalle et al. 2018; Bailey et al. 2018; Joshi et al. 2020). Interestingly, ERBB2 and EGFR are known to be associated with Erlotinib (one of the drugs we studied) that demonstrated cross-tissue generalization to non-solid tissues. EGFR activation drives the development of most head and neck tumors, and in a large proportion of glioblastoma and lung adenocarcinoma cases (Thomas and Weihua 2019). More recently, overexpression of EGFR was reported in a subset of acute myeloid leukemia patients with poor prognosis (Nath et al. 2020). A related receptor tyrosine kinase ERBB2 is frequently activated in breast cancer and, more rarely, in several other tumor types (Iqbal and Iqbal 2014). ERBB2 amplification and activation are associated with poor prognosis and the resistance to anti-EGFR treatments (Yonesaka et al. 2011), including Erlotinib (Goss et al. 2018). Later, putative driver ERBB2 mutations were found in three leukemia patients (Joshi et al. 2020). Their effect on proliferation rate and sensitivity to ERBB inhibitors was shown in cellular assays ( Joshi et al. 2020).

We would like to note that the promising Velodrome results should be interpreted with caution due to the lack of drug response values in our TCGA association analyses. Therefore, direct experimental validation of cross-tissue generalization and Velodrome predictions is a direction of future research.

## Conclusion

In this paper, we proposed Velodrome, a transfer learning method for drug response prediction based on gene expression data. Velodrome is the first semi-supervised method of out-of-distribution generalization. We trained Velodrome on cell line datasets with drug response (measured in AAC) and patient datasets without drug response (i.e., unlabelled data) as source domains and successfully validated it on different target domains such as cell lines, PDX samples, and patient data across three chemotherapy agents and one targeted therapeutic. Our results suggest that Velodrome outperforms state-of-the-art methods of drug response prediction and transfer learning in terms of Pearson and Spearman correlations (on cell lines) and in terms of AUROC and AUPR (on PDX samples and patients). Moreover, we analyzed the biological significance of the predictions made by Velodrome and provided substantial evidence that these predictions have statistically significant associations with the expression level of numerous known target genes of the studied drugs in a tissue well-represented in our source domains, i.e. kidney cancer, and a tissue under-represented in our source domains, i.e. prostate cancer. Finally, we also demonstrated that Velodrome generalizes to new tissue types that were completely absent in the source domains. All these results demonstrate the superior out-of-distribution generalization capability of the Velodrome model and suggest that Velodrome may guide pharmacogenomics and precision oncology more accurately.

## Methods

### Data Preprocessing

We obtained all cell line datasets from the ORCESTRA platform (Mammoliti et al. 2020) which stores pharmacogenomics datasets in PharmacoSet (PSet) *R* objects. Samples with missing values were removed from both the gene expression and drug response data. The cell line datasets were generated via the same drug screening assay (CellTiter Glo) preprocessed using the PharmacoGx package version 2.0.5 (Smirnov et al. 2016) and are also comparable in terms of gene expression data which was preprocessed via Kallisto_0.46.1 (Bray et al. 2016). We also removed all the cell lines originating from non-solid tissue types from the cell line datasets.

We obtained the TCGA dataset via the Firehose (http://gdac.broadinstitute.org/) 28.01.2016. Expression values were converted to Transcripts Per Million (TPM) and log2-transformed. The PDX and clinical trial datasets were preprocessed similar to the approach described in (Hossein Sharifi-Noghabi et al. 2019). For Docetaxel and Paclitaxel patient data, the accession code is GSE25065 and for Erlotinib, the accession code is GSE33072.

For all of the employed datasets, all gene names were mapped to Entrez gene ids and the expression data were obtained before treatment and the response outcome after treatment. We reduced the number of genes to 2128 genes obtained from (Manica et al. 2019). After the preprocessing, all of the available datasets for each drug had the same number of genes (Supplementary Table 1).

### The Velodrome Method

We propose Velodrome, a method of drug response prediction using labeled and unlabeled source domains to build a predictive model that generalizes to unseen domains. To achieve this goal, Velodrome requires three different characteristics: 1) being predictive of drug response, 2) being generalizable to unseen domains, and 3) achieving these goals by taking both labeled and unlabeled data. We designed the objective function of Velodrome to meet these requirements by combining three loss functions: 1) a standard supervised loss based on labeled data to ensure that the model is predictive, 2) an alignment loss to ensure that the model has generalization capabilities, and 3) a consistency loss that exploits the unlabeled data.

#### Problem Definition

Following the notation of (Pan and Yang 2010), a domain *D* is defined by a raw input space **X**, a probability distribution *p*(*X*)and a corresponding dataset *X* = {*x*_1_, *x*_2_,…, *x*_*n*_} with *x*_*i*_ ∈**X**. A task 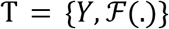}is associated with *D* = {*X*, *p*(*X*)}and is defined by a label space *Y* ∈**Y** and a predictive function 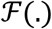 which is learned from training data (*X, Y*) ∈**X**×**Y**. In our case, **Y**∈ [0, 1], which makes drug response prediction a regression problem.

Given multiple labeled and unlabeled source domains denoted by 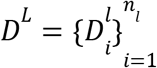 and 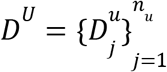, the goal is to learn the predictive function 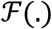 which is implemented through a neural network.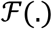consists of a shared (across all source domains) feature extractor *F*_θ_(*X*) parameterized by θ, which maps *X* to latent features *Z*, and domain-specific 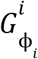 predictors parametrized by ϕ_*i*_, which takes *Z*_*i*_ (the extracted features of*D*_*i*_) as input and makes predictions (of the drug response) 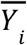 for this source domain. θ and 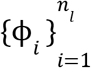 are being optimized using an objective function 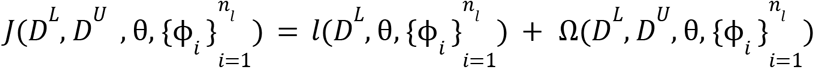 with a supervised loss*l*(.) and some regularization terms Ω(.).

In drug response prediction, we have access to labeled source domains such as cell line datasets and unlabeled source domains such as cancer patients in TCGA. The goal is to learn a model that makes accurate predictions on patients, PDXes, or other cell lines as target domains that it may see during deployment. This is similar to out-of-distribution generalization (also known as domain generalization), where the goal is to optimize parameters of the model (θand 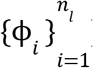) in order to make the model generalizable and predictive of unseen domains. Out-of-distribution generalization assumes that there exists a *d*-dimensional latent feature space *Z* ∈ *R*^*d*^ that is invariant, predictive, and generalizable to seen and unseen domains on this given space.

#### Shared feature extractor

To map the raw input gene expression data to the latent space, Velodrome utilizes a feature extractor which is shared across all labeled and unlabeled source domains:

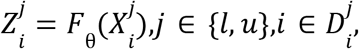

where, 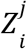 denotes the features extracted by the feature extractor*F*_θ_ (.)from 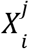, the samples obtained from the *i*-th domain of type *j* (labeled or unlabeled). These extracted (latent) features will be provided as input to the domain-specific predictors.

#### Domain-specific predictors

To make predictions for the samples in the source domains, Velodrome utilizes *n*_*l*_ domain-specific predictors, meaning the number of domain-specific predictors that Velodrome utilizes is the same as the number of labeled source domains. These predictors are formulated as follows:

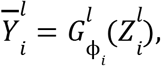

where, 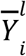denotes the predictions for the *i*-th labeled source domain obtained from predictor 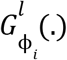associated with the *i*-th labeled source domain and parameterized by ϕ_*i*_. These predictions will be utilized to optimize the parameters of the feature extractor and the *i*-th predictor.

#### Supervised loss

To make the extracted latent features predictive of the drug response, Velodrome utilizes a standard supervised loss as follows:

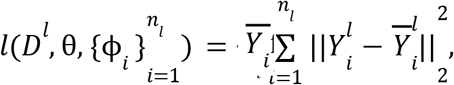

where, *l*(.)denotes a standard supervised loss function in the form of the mean squared error (MSE). It is important to note that the parameters of the feature extractor are optimized by the total supervised loss but the parameters of the *i*-th predictor are optimized only by the supervised loss on predictions of the *i*-th predictor.

#### Alignment loss

Optimizing the parameters of the Velodrome model using only the supervised loss is likely to lead to overfitting to the labeled source domains. Therefore, we need an additional loss function to avoid overfitting to the source domains and to make the latent representation generalizable to unseen domains. To achieve this, Velodrome utilizes the CORAL loss function that regularizes the covariance matrices across input domains and has demonstrated state-of-the-art performance for learning invariant representations in computer vision applications (Sun and Saenko 2016; Gulrajani and Lopez-Paz 2020). The CORAL loss is defined as follows:

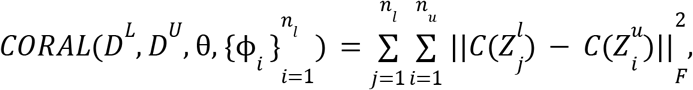

where, C(.) is the covariance operator which receives the extracted features of a source domain and returns the covariance matrix of those features as follows:

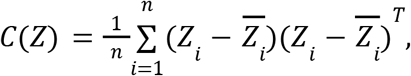

where, *n* is the number of samples and 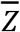denotes the mean vector. Regularizing the covariance matrices across source domains ensures learning invariant feature vectors. It is important to note that the objective function of Velodrome requires a combination of supervised and alignment loss because optimizing only the alignment loss is likely to lead to a trivial “zero” solution where all domains are mapped to the same point (Sun and Saenko 2016).

#### Consistency loss

Aligning the extracted features of the different domains imposes a strict constraint on learning an invariant latent representation because it disregards the unique domain-specific aspects of different source domains. To alleviate this, Velodrome utilizes a consistency loss to ensure that it learns a hypothesis invariant representation, i.e. predictions across source domains are similar when using different predictors. For example, if we have two predictors 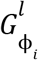 and 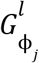, we want them to generate similar predictions for the same unlabeled source domain. This consistency loss is defined as follows:

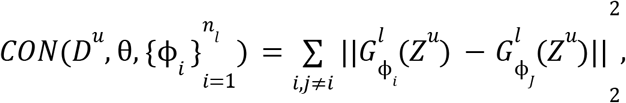

where, *Z*^*u*^ are extracted features for samples in a given unlabeled source domain and *MSE* is the mean squared error.

#### Objective function

Putting all of the loss functions together, the objective function of Velodrome is as follows:

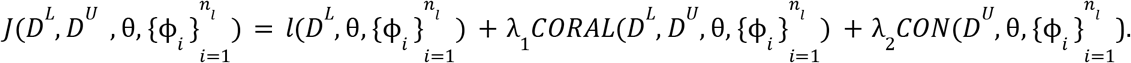

where, λ_1_ and λ_2_ = 1 − λ_1_ denote the regularization coefficients for the coral loss and consistency loss, respectively. Therefore, the function Ω(.)that we defined in the problem definition is given byΩ(.) = λ_1_ *CORAL*(.) + λ_2_ *CON*(.). λ_1_ and λ_2_ control balancing between learning a domain-invariant representation and learning a hypothesis-invariant representation because the alignment loss ensures learning a domain-invariant representation, while the consistency loss ensures learning a hypothesis-invariant representation. The training steps of the Velodrome method are presented in Algorithm 1.

**Algorithm 1:**
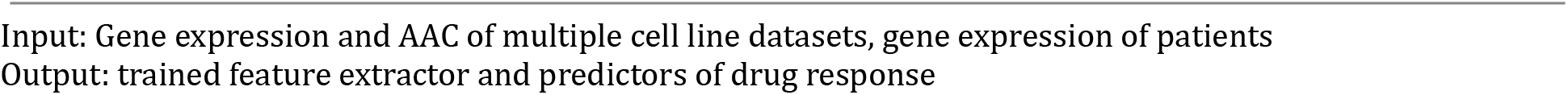
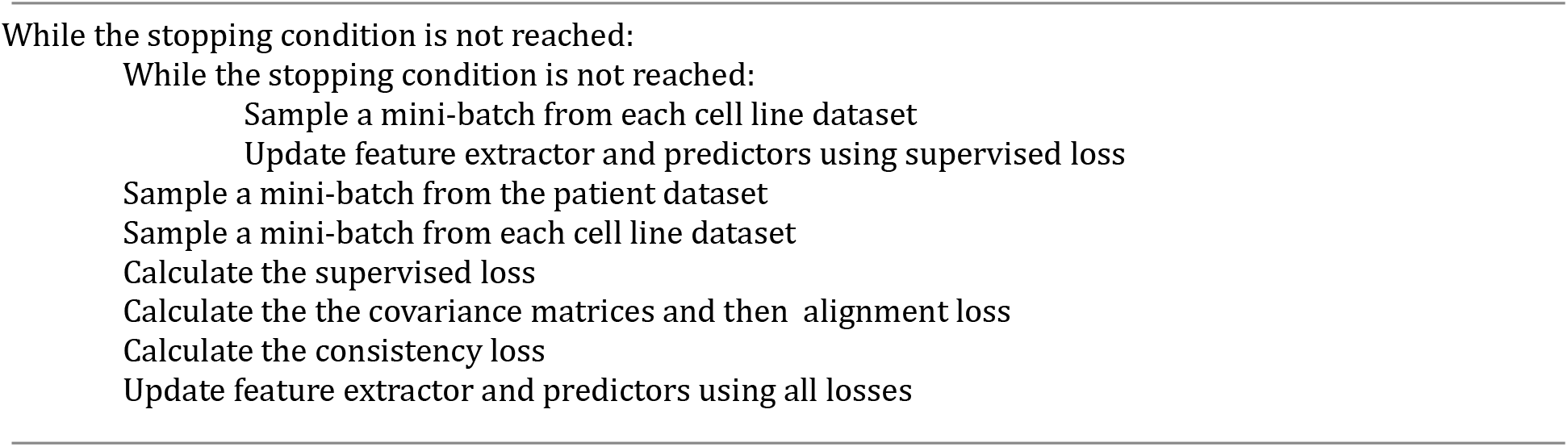
Velodrome.

#### Velodrome at test time

For a target sample *x*^*t*^, elodrome makes prediction as follows:

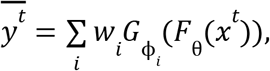

where, *w*_*i*_ denotes the average supervised loss for the predictions of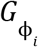 normalized via a softmax function such that 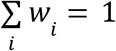.This means the final prediction will be a result of a weighted average of all predictors, and more accurate predictors will have higher weights.

### Implementation Detail

#### Hyper-parameters

We considered a wide range of values for each hyper-parameter of the Velodrome model and optimized these values via a random search separately for each drug. The sets of values considered are as follows:

*Epoch* = [10, 50, 100, 200], *Learning rate* (*LR*) = [0. 0001, 0. 001, 0. 01, 0. 0005, 0. 005, 0. 05]

*Dropout* (*DR*) = [0. 1, 0. 3, 0. 5, 0. 8]

*Weight Decay* (*WD*) = [0. 001, 0. 0001, 0. 01, 0. 05, 0. 005, 0. 0005]

λ_1_ = [1, 0. 1, 0. 2, 0. 3, 0. 4, 0. 5, 0. 01, 0. 05, 0. 001, 0. 005, 0. 0001, 0. 0005]

*Mini batch size* (*MB*) = [17, 33, 65, 129]

We considered separate learning rates and weight decays for the feature extractor and each predictor, but they all used the same sets of possible values.

We split the labeled cell line datasets (CTRPv2 and GDSCv2) into train and validation and considered 90% for train and 10% for validation. We merged the train splits into one training dataset and similarly, merged the validation splits into one validation set and used the merged validation set to optimize the values of these hyper-parameters.

#### Velodrome architecture

We followed previous works and designed predefined architectures (denoted by HD) for Velodrome (Noghabi et al. 2021; Sakellaropoulos et al. 2019). For the feature extractor, the first architecture has two hidden layers with the size 512 × 128, the second one has two layers with the size 256 × 256, the third one has three hidden layers with the size 128 × 128 × 128 and the last architecture has four hidden layers with the size 64 × 64 × 64 × 64. We considered a batch normalization layer followed by an activation function (which we considered the Relu, the Tanh, and Sigmoid functions) as well as a dropout after the activation function for each hidden layer. The predictors have only one layer *HD* × 1, where HD denotes the size of the last layer in the feature extractor. The final hyper-parameter and architecture of Velodrome for the studied drugs are as follows:

Drug: Epoch, MB, DR, WD1, WD2, WD3, HD, LR1, LR2, LR3, λ_1_,λ_2_

Docetaxel: 10, 65, 0.1, 0.05, 0.0005, 0.0001, 3, 0.001, 0.005, 0.0005, 0.2, 0.8

Gemcitabine: 10, 17, 0.1, 0.0001, 0.005, 0.01, 2, 0.01, 0.005, 0.05, 0.005, 0.99

Erlotinib: 50, 129, 0.1, 0.05, 0.005, 0.0005, 2, 0.001, 0.01, 0.001, 0.01, 0.99

Paclitaxel: 50, 129, 0.1, 0.005, 0.05, 0.005, 2, 0.05, 0.0005, 0.0001, 0.3, 0.7

WD1, WD2, and WD3 refers to the values we used for the feature extractor, predictor 1, and predictor 2, respectively (similar for LR1, LR2, and LR3).

For re-running and the ablation study of the trained models, we considered these random values for the random seed:

*Seed* = [1, 21, 42, 84, 168, 336, 672, 1344, 2688, 5376].

We used 42 for the majority of the analyses in the paper (because it’s the answer to life, the universe and everything!).

We used the same ranges for all of the baseline methods whenever using those values was applicable. For DeepAll-ERM and Ridge-ERM we used the existing implementations here: (https://github.com/bhklab/PGx_Guidelines), for PRECISE, we used the existing implementations here: (https://github.com/NKI-CCB/PRECISE). For Mean Teacher, we adopted an existing implementation for computer vision and modified it for this problem here: (https://github.com/CuriousAI/mean-teacher).

All of the deep neural network implementations were in the PyTorch framework and we employed the Adagrad optimizer to optimize the parameters of Velodrome as well as the baselines wherever applicable.

#### Performance evaluation

We employed the Scikit-learn and Scipy Python packages for the evaluation purposes. To be more specific, we utilized scikit-learn to calculate the AUROC and AUPR (for PDX samples and Patients) and we utilized the Scipy to calculate Pearson and Spearman correlations (for cell lines). For the association study, we utilized statsmodels.api Python package to fit the multiple linear regression and obtain the P-values and we obtained the list of known associated target genes for each drug by querying the PharmacoDB resource (Smirnov et al. 2018). We did not perform statistical tests on the performance of Velodrome compared to the other methods to evaluate level of significance. For the association studies of well-/under-represented tissues types, we performed a two-tailed t-test on the regression coefficients with the null hypothesis that a given coefficient (corresponding to a gene) is zero and has no significant association with drug response. For the correlation analysis on the unseen tissues (non-solid tissue types), we employed a two-tailed test, testing the specific null hypothesis that the population correlation is 0 against a two-tailed alternative.

## Acknowledgement

We would like to thank Hossein Asghari (Ocean Genomics) and Shuman Peng (Simon Fraser University) for their support. We also would like to thank the Vancouver Prostate Centre and Compute Canada (West Grid) for providing the computational resources for this research.

## Funding

This work was supported by a Discovery Grant from the National Science and Engineering Research Council of Canada (to M.E.), Canada Foundation for Innovation (33440 to C.C.C.), The Canadian Institutes of Health Research (PJT-153073 to C.C.C.), Terry Fox Foundation (201012TFF to C.C.C.), and The Terry Fox New Frontiers Program Project Grants (1062 to C.C.C.).

## Authors’ contributions

Study concept and design: H.S-N., M.E.

Deep learning design, implementations, and analysis: H.S-N.

Data preprocessing, analysis, and interpretation: H.S-N., O.Z.

Experiments: H.S-N., P.AH.

Analysis and interpretation of results: H.S-N., P.AH., O.Z.

Supervision: C.C.C., M.E.

## Conflict of Interest

None declared.

## Data availability statement

All the final preprocessed data employed in this paper are publicly available here: https://zenodo.org/record/4793442#.YK1HVqhKiUk

All the raw data before preprocessing are also publicly available as follows:

1. Cell line datasets with gene expression and drug response data including, CTRPv2, GDSCv2, and gCSI were downloaded from the ORCESTRA platform (Mammoliti et al. 2020)
2. TCGA cohorts with gene expression data were downloaded from the Firehose (http://gdac.broadinstitute.org/) 28.01.2016. Drug response data for TCGA cohorts was obtained from (Ding, Zu, and Gu 2016).
3. PDX datasets (gene expression with drug response data) were obtained from the supplementary material of (Gao et al. 2015).
4. Patient dataset (gene expression with drug response data) for Docetaxel and Paclitaxel were obtained from the accession code GSE25065 and for Erlotinib from the accession code of GSE33072.

## Code availability statement

All the codes, model objects, and supplementary material to run and reproduce our experimental results are publicly available here: https://github.com/hosseinshn/Velodrome DOI: https://doi.org/10.5281/zenodo.5164625

We also provided a conda environment to ensure version compatibility for future users.

## Supplementary information

**Supplementary Table 1.** Characteristics of the employed datasets

| Dataset | Drug | Type | Domain | Label | Tissue | #Samples | #Genes |
| --- | --- | --- | --- | --- | --- | --- | --- |
| CTRPv2 | Docetaxel | Cell line | Source | AAC | Solid | 292 | 1453 |
| GDSCv2 | Docetaxel | Cell line | Source | AAC | Solid | 234 | 1453 |
| TCGA-LUAD | Docetaxel | Patient | Source | Unlabeled | Solid | 507 | 1453 |
| TCGA-BRCA | Docetaxel | Patient | Source | Unlabeled | Solid | 1051 | 1453 |
| TCGA-PAAD | Docetaxel | Patient | Source | Unlabeled | Solid | 131 | 1453 |
| gCSI | Docetaxel | Cell line | Target | AAC | Solid | 280 | 1453 |
| GSE25065D | Docetaxel | Patient | Target | RECIST | Solid | 51 | 1453 |
| CTRPv2 | Gemcitabine | Cell line | Source | AAC | Solid | 514 | 2080 |
| GDSCv2 | Gemcitabine | Cell line | Source | AAC | Solid | 226 | 2080 |
| TCGA-LUAD | Gemcitabine | Patient | Source | Unlabeled | Solid | 507 | 2080 |
| TCGA-BRCA | Gemcitabine | Patient | Source | Unlabeled | Solid | 1051 | 2080 |
| TCGA-PAAD | Gemcitabine | Patient | Source | Unlabeled | Solid | 131 | 2080 |
| gCSI | Gemcitabine | Cell line | Target | AAC | Solid | 277 | 2080 |
| TCGA-Gem | Gemcitabine | Patient | Target | RECIST | Solid | 66 | 2080 |
| PDXE | Gemcitabine | PDX | Target | RECIST | Solid | 25 | 2080 |
| CTRPv2 | Erlotinib | Cell line | Source | AAC | Solid | 607 | 2066 |
| GDSCv2 | Erlotinib | Cell line | Source | AAC | Solid | 230 | 2066 |
| TCGA-LUAD | Erlotinib | Patient | Source | Unlabeled | Solid | 507 | 2066 |
| TCGA-BRCA | Erlotinib | Patient | Source | Unlabeled | Solid | 1051 | 2066 |
| TCGA-PAAD | Erlotinib | Patient | Source | Unlabeled | Solid | 131 | 2066 |
| gCSI | Erlotinib | Cell line | Target | AAC | Solid | 283 | 2066 |
| GSE33072 | Erlotinib | Patient | Target | RECIST | Solid | 25 | 2066 |
| PDXE | Erlotinib | PDX | Target | RECIST | Solid | 21 | 2066 |
| CTRPv2 | Paclitaxel | Cell line | Source | AAC | Solid | 445 | 1452 |
| GDSCv2 | Paclitaxel | Cell line | Source | AAC | Solid | 230 | 1452 |
| TCGA-LUAD | Paclitaxel | Patient | Source | Unlabeled | Solid | 507 | 1452 |
| TCGA-BRCA | Paclitaxel | Patient | Source | Unlabeled | Solid | 1051 | 1452 |
| TCGA-PAAD | Paclitaxel | Patient | Source | Unlabeled | Solid | 131 | 1452 |
| gCSI | Paclitaxel | Cell line | Target | AAC | Solid | 284 | 1452 |
| GSE25065P | Paclitaxel | Patient | Target | RECIST | Solid | 84 | 1452 |
| PDXE | Paclitaxel | PDX | Target | RECIST | Solid | 43 | 1452 |
| TCGA-PRAD | Docetaxel | Patient | Target | Unlabeled | Solid | 498 | 1453 |
| TCGA-KIRC | Docetaxel | Patient | Target | Unlabeled | Solid | 534 | 1453 |
| TCGA-PRAD | Paclitaxel | Patient | Target | Unlabeled | Solid | 498 | 1452 |
| TCGA-KIRC | Paclitaxel | Patient | Target | Unlabeled | Solid | 534 | 1452 |
| TCGA-PRAD | Gemcitabine | Patient | Target | Unlabeled | Solid | 498 | 2080 |
| TCGA-KIRC | Gemcitabine | Patient | Target | Unlabeled | Solid | 534 | 2080 |
| TCGA-PRAD | Erlotinib | Patient | Target | Unlabeled | Solid | 498 | 2066 |
| TCGA-KIRC | Erlotinib | Patient | Target | Unlabeled | Solid | 534 | 2066 |
| CTRPv2 | Docetaxel | Cell line | Source | AAC | Non-solid | 62 | 1453 |
| GDSCv2 | Docetaxel | Cell line | Source | AAC | Non-solid | 69 | 1453 |
| gCSI | Docetaxel | Cell line | Target | AAC | Non-solid | 50 | 1453 |
| CTRPv2 | Paclitaxel | Cell line | Source | AAC | Non-solid | 100 | 1452 |
| GDSCv2 | Paclitaxel | Cell line | Source | AAC | Non-solid | 67 | 1452 |
| gCSI | Paclitaxel | Cell line | Target | AAC | Non-solid | 50 | 1452 |
| CTRPv2 | Gemcitabine | Cell line | Source | AAC | Non-solid | 129 | 2080 |
| GDSCv2 | Gemcitabine | Cell line | Source | AAC | Non-solid | 69 | 2080 |
| gCSI | Gemcitabine | Cell line | Target | AAC | Non-solid | 50 | 2080 |
| CTRPv2 | Erlotinib | Cell line | Source | AAC | Non-solid | 135 | 2066 |
| GDSCv2 | Erlotinib | Cell line | Source | AAC | Non-solid | 68 | 2066 |
| gCSI | Erlotinib | Cell line | Target | AAC | Non-solid | 49 | 2066 |
| TCGA-LAML | Docetaxel | Patient | Target | Unlabeled | Non-solid | 173 | 1443 |
| TCGA-DLBC | Docetaxel | Patient | Target | Unlabeled | Non-solid | 48 | 1443 |
| TCGA-LAML | Erlotinib | Patient | Target | Unlabeled | Non-solid | 173 | 2047 |
| TCGA-DLBC | Erlotinib | Patient | Target | Unlabeled | Non-solid | 48 | 2047 |
| TCGA-LAML | Gemcitabine | Patient | Target | Unlabeled | Non-solid | 173 | 2061 |
| TCGA-DLBC | Gemcitabine | Patient | Target | Unlabeled | Non-solid | 48 | 2061 |
| TCGA-LAML | Paclitaxel | Patient | Target | Unlabeled | Non-solid | 173 | 1442 |
| TCGA-DLBC | Paclitaxel | Patient | Target | Unlabeled | Non-solid | 48 | 1442 |

**Supplementary Table 2.** Characteristics of the employed cell line datasets in terms of drug response data

| Drug | Dataset | Type | Mean | sd |
| --- | --- | --- | --- | --- |
| Docetaxel | Train | Cell line | 0.36 | 0.24 |
| Docetaxel | Test | Cell line | 0.3 | 0.16 |
| Gemcitabine | Train | Cell line | 0.36 | 0.24 |
| Gemcitabine | Test | Cell line | 0.3 | 0.13 |
| Erlotinib | Train | Cell line | 0.12 | 0.1 |
| Erlotinib | Test | Cell line | 0.12 | 0.1 |
| Paclitaxel | Train | Cell line | 0.33 | 0.28 |
| Paclitaxel | Test | Cell line | 0.41 | 0.15 |

Train indicates CTRPv2 and GDSCv2 cell line datasets together.

**Supplementary Table 3.** Characteristics of the employed patient/PDX datasets in terms of drug response data

| Drug | Dataset | Type | #Sensitive | #Resistant |
| --- | --- | --- | --- | --- |
| Docetaxel | Test | Patient | 46 | 5 |
| Gemcitabine | Test | PDX | 7 | 18 |
| Gemcitabine | Test | Patient | 25 | 41 |
| Erlotinib | Test | PDX | 3 | 18 |
| Erlotinib | Test | Patient | 13 | 12 |
| Paclitaxel | Test | PDX | 5 | 38 |
| Paclitaxel | Test | Patient | 58 | 26 |

**Supplementary Figure 1.**
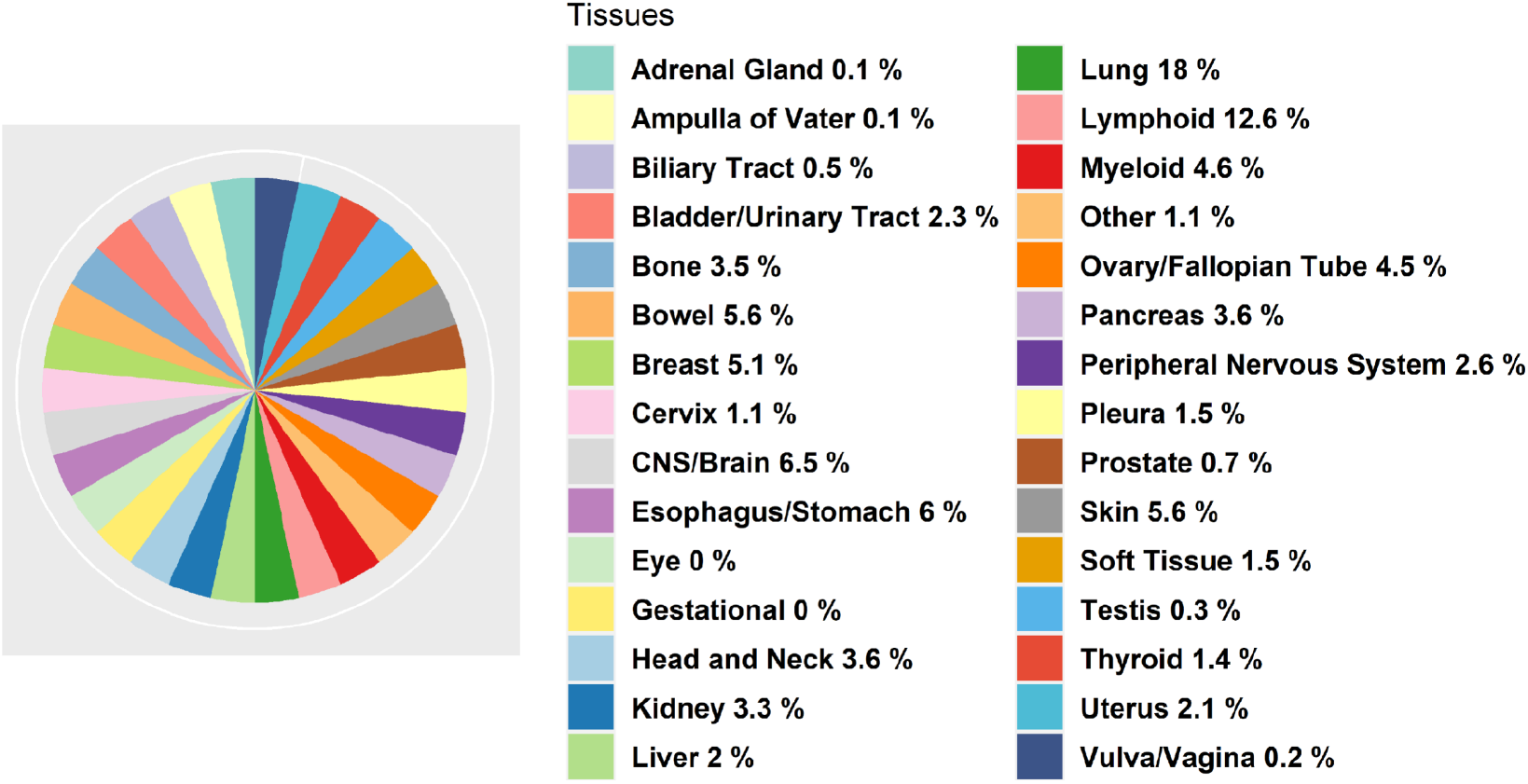
The percentage of tissue types in CTRPv2 and GDSCv2 cell line datasets combined.

## Notes

https://github.com/hosseinshn/Velodrome

### Competing Interest Statement

The authors have declared no competing interest.

### Summary of Updates

Substantial revision for cross-tissue generalization

